# Competition between chemoattractants causes unexpected complexity and can explain negative chemotaxis

**DOI:** 10.1101/2022.12.07.519354

**Authors:** Adam Dowdell, Peggy Paschke, Peter Thomason, Luke Tweedy, Robert H. Insall

## Abstract

Negative chemotaxis, where eukaryotic cells migrate away from repellents, is important throughout biology, for example in nervous system patterning and resolution of inflammation. However, the mechanisms by which molecules repel migrating cells are unknown. Here, we use a combination of modelling and experiments with Dictyostelium cells to show that competition between different ligands that bind to the same receptor leads to effective chemorepulsion. 8-CPT-cAMP, widely described as a simple chemorepellent, is inactive on its own, and only repels cells if it interacts with the attractant cAMP. If cells degrade either competing ligand, the pattern of migration becomes more complex; cells may be repelled in one part of a gradient but attracted elsewhere, leading to populations moving in different directions in the same assay, or converging in an arbitrary place. More counterintuitively still, two chemicals can each attract cells on their own, but repel cells when combined together. We have thus identified a new mechanism that drives reverse chemotaxis, verified by mathematical models and experiments with real cells, and important anywhere several ligands compete for the same receptors.

## Introduction

Negative chemotaxis (also called chemorepulsion(Tharp et al., 2006)) occurs when cells are exposed to a spatial gradient of a signalling molecule and move down-gradient, towards the region with the lowest concentration. It is thus the opposite of normal (“positive”) chemotaxis, in which cells move up-gradient, towards the highest concentration.

Negative chemotaxis is implicated in a wide range of whole-organism processes including development (Pasquale, 2000), immunity (Colom et al., 2015, Consalvo et al., 2022) pathogenesis (Zaki et al., 2006) and cancer metastasis (Jongsma et al., 2011). For example, neurites in the developing brain avoid areas high in semaphorins (Sakai and Halloran, 2006), and semaphorin knockout mice die at or before birth (Fiore et al., 2005). Negative chemotaxis is also important for normal immune cell behaviour. Neutrophils are attracted to infections and sites of tissue damage by chemoattraction, but when the damage has been resolved they can migrate in reverse, back into the bloodstream (Holmes et al., 2012). This is thought to be mediated by negative chemotaxis through G-protein coupled receptors (GPCRs), often using chemokines and complement fragments. Notably, these molecules normally act positively, promoting attraction rather than repulsion. These multiple observations of GPCR-mediated repulsion remain unexplained.

In bacteria, negative chemotaxis is widespread (Tso and Adler, 1974). It works because receptors are tuned by protein methylation to register a “background” concentration of attractant or repellent (Toews et al., 1979). If the attractant concentration is higher than the background, the receptor mediates an attractive response; if lower, the response is negative. Thus one receptor can communicate both positive and negative chemotactic stimuli. Likewise, Dictyostelium phototaxis works negatively (Bonner et al., 1988, Wilkins et al., 2000); ammonia repels the tip of migrating slugs, and light promotes ammonia release on the far side of the light source.

By comparison, eukaryotic chemorepulsion is difficult to explain. Eukaryotic chemotaxis integrates temporal information with spatial data (Gerisch and Keller, 1981), where the cell compares receptors at different places in the cell. The resulting spatial information is used to steer the cell (Devreotes and Zigmond, 1988). Much research has addressed the mechanisms by which attractive receptor stimuli reinforce actin protrusions. The GPCRs that are used in chemotaxis have a wide range of ligand specificities. These varied inputs are integrated at the level of G-protein coupling. When G-proteins are activated, their βγ-subunits positively reinforce local actin polymerisation (Neptune et al., 1999, Neptune and Bourne, 1997), connecting signals from outside the cell to the intracellular processes that control cell movement.

It is difficult to envisage how negative stimuli can be integrated into this type of chemotaxis pathway. G protein βγ-subunits are not thought to be specific, and there is little to suggest that positive and negative stimuli act through different βγ-subunits. Furthermore, when negative chemotaxis has been analysed in detail, the same receptors promote both positive and negative effects (Malet-Engra et al., 2015, Tharp et al., 2006). This makes it hard to understand how G-proteins could mediate negative steering. Many eukaryotic cells simultaneously use a wide range of receptors to different attractants; neutrophils, for example, express hundreds of distinct receptors. This makes the prokaryotic scheme, in which repellents act by decreasing the number of activated molecules of one class of receptor, unfeasible; if a large number of different receptors each have a background activity, the overall effect of depressing one receptor type is small.

Despite this, negative chemotaxis clearly occurs, and is important in normal physiology (Nourshargh et al., 2016, Robertson et al., 2014). This paper examines the mechanisms that can allow this.

Recent work has focussed on cell guidance by self-generated attractant gradients (Dalle Nogare and Chitnis, 2017, Dona et al., 2013, Insall et al., 1992, Insall et al., 2022, Muinonen-Martin et al., 2014, Shellard and Mayor, 2021, Tweedy et al., 2016, Tweedy and Insall, 2020, Venkiteswaran et al., 2013). These are exceptionally dynamic, and the evolving ligand profiles are near-impossible to measure directly (Tweedy et al., 2016). In contrast, the behaviour of cells is readily observable and can be accurately predicted using computational models (Neilson et al., 2011). These models need not include complex receptor behaviour (such as recycling, multimerization and adaptation) to show strong predictive power. This reveals that the key determinant of cell behaviour is the profile of activated receptors. Here, we test the idea that different ligands can make complex and negative gradients by competing for simple, positively-acting receptors. We combine computational modelling with experiments to show that molecules described as chemorepellents can work by interfering with positive signals, rather than via undiscovered negative signalling pathways.

## Results

### Chemorepellents work by competing with positive agonist signals

8-CPT-cAMP has been described as a strong chemorepellent for Dictyostelium (Cramer et al., 2018, Keizer-Gunnink et al., 2007). As an analogue of cAMP, 8-CPT-cAMP is principally detected through the cAR1 receptor, which normally mediates a strongly attractive response to cAMP(Pupillo et al., 1992). Since it is difficult to imagine this receptor acting both positively and negatively, we tested a different hypothesis - that it was acting as a receptor antagonist rather than a direct repellent. If 8-CPT-cAMP acts as a competitive inhibitor, binding to the receptor but not activating it, it will not stimulate G-proteins and will give no directional cues on its own. In the presence of a stimulatory ligand, however, the 8-CPT-cAMP would compete for receptors; higher concentrations of 8-CPT-cAMP would keep more receptors in an inactive state. A gradient of 8-CPT-cAMP could therefore interact with uniform cAMP to create a reverse gradient of activated cAR1 receptors (represented in Fig. 1A).

**Fig. 1:**
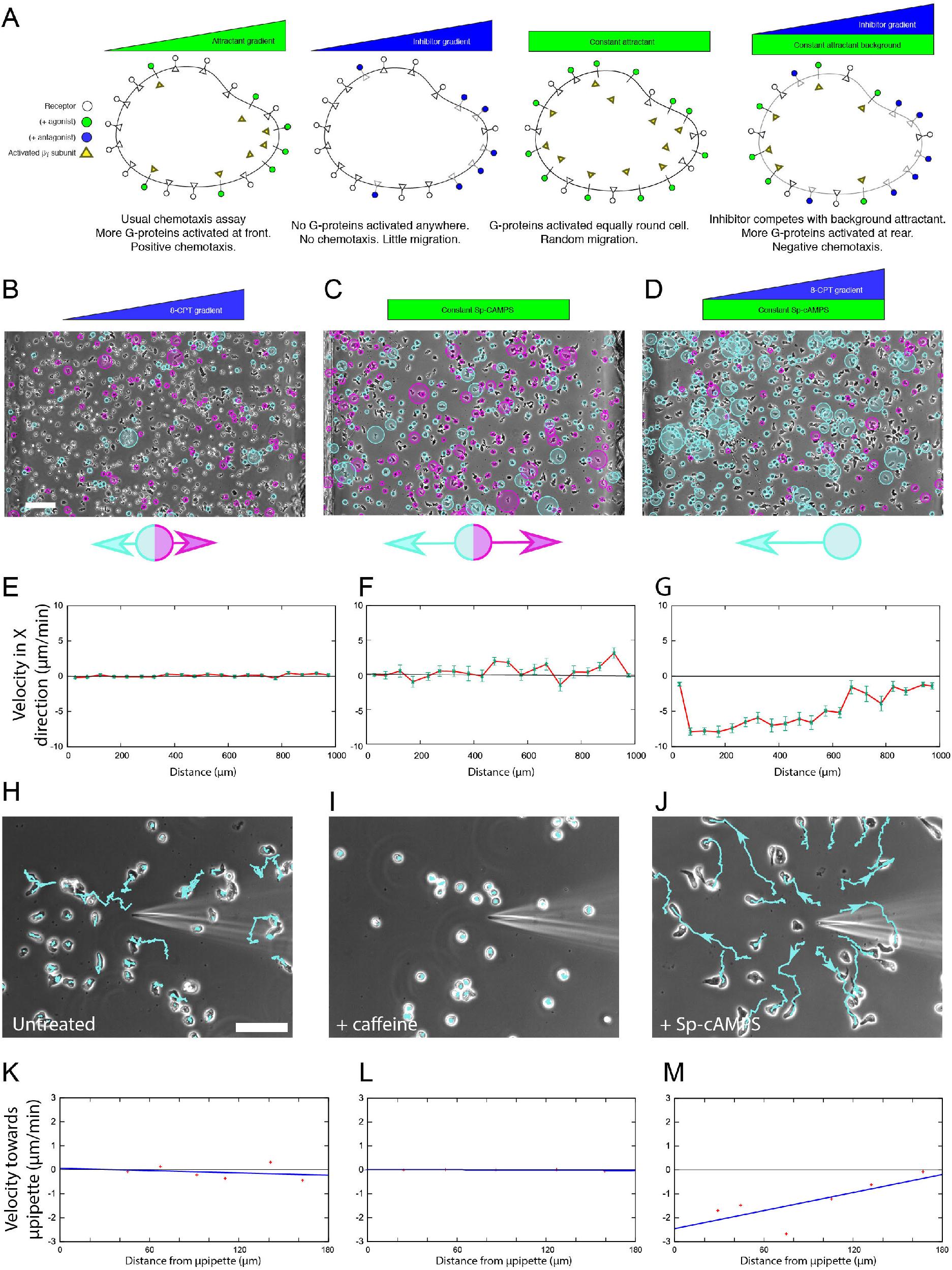
Repulsion by 8-CPT-cAMP depends on the presence of a competing agonist. **(A)** Diagram of guidance by receptor competition. Agonists are attractive because a concentration gradient creates a gradient of activated receptors and G-proteins (far left). A similar gradient of antagonist occupies receptors similarly, but does not activate receptors or G-proteins (left). Homogeneous attractant (right) activates receptors and G-proteins, but there is no directional bias so the cells are not steered. A combination of homogeneous agonist and a gradient of antagonist causes a steering in the opposite orientation of the gradient (far right). **(B-D)** Chemotaxis chamber assays exposing cells to (B) 0-200μM 8-CPT-cAMP, (C) uniform 2μM Sp-cAMPS, (D) both. Tracking software highlights cell movement to the left with a blue circle and to the right with a purple circle. Larger circles indicate a higher velocity component in the specified direction. Scale bar 100μm. **(E-F)** X-direction (i.e. rightward) velocity across a minimum of 3 repeats of the experiments in (B-D). 8-CPT-cAMP alone causes almost no movement (E). Uniform Sp-cAMPS drives some chemokinetic stimulus, but no chemotaxis (F). Substantial movement away from high concentrations of 8-CPT-cAMP is apparent when both are included (G). **(H-J)** NC4 cells stimulated by 10mM 8-CPT-cAMP from a micropipette without caffeine (H), with caffeine (I), and with both caffeine and agonist background (2μM Sp-cAMPS; J). Scale bar 40μm. **(K-M)** Radial velocity toward the micropipette as a function of distance from it (K) without caffeine, (L) with caffeine, (M) with both caffeine and 2μM Sp-cAMPS. Repulsion is at best slight (K), with caffeine treatment erasing it completely (L). The introduction of an agonist makes the effect strong, and strongest close to the needle (M). Points show mean ± SE of technical replicates for cells at the indicated positions in a representative experiment (n≥30, 48 and 28 respectively).

We tested this prediction using direct-view chemotaxis chambers(Muinonen-Martin et al., 2010), which generate a defined gradient between two reservoirs (Fig. 1 B-D; Movie 2). Gradients of 8-CPT-cAMP do not give a steering response on their own (Fig. 1B, measured in Fig. 1E), but strongly repel cells in the presence (Fig. 1D, G) of homogeneous levels of a non-degradeable cAMP analogue, Sp-cAMPS, which strongly activates cAR1 but is not degraded(Theibert et al., 1986). Sp-cAMPS on its own gave no directional migration (Fig. 1C, F).

Thus the 8-CPT-cAMP does not repel cells directly, but works very effectively through competition with an active ligand. This is not negative chemotaxis as usually understood - cells do not respond directly to the repellent - but it is remarkably effective (Movie 2).

This implied that earlier results, in which 8-CPT-cAMP was found as a direct chemorepellent (Cramer et al., 2018), worked through interaction with endogenous attractants. *Dictyostelium* cells complicate experimental results by secreting their own cAMP, which can be blocked using caffeine(Theibert and Devreotes, 1983). When we examined the chemotaxis of untreated cells using a micropipette, we found that wild type cells were weakly repelled by 8-CPT-cAMP (Fig. 1H, K), but we could never detect repulsion of caffeine-treated cells. (Fig. 1I, L). This suggests that cAMP produced by normal cells is essential for the observed repulsion. We therefore tested the effect of adding a homogeneous background of Sp-cAMPS. This fully rescued chemorepulsion, causing cells to migrate directly and efficiently away from 8-CPT-cAMP released from a micropipette (Fig 1J, M; Movie 1).

Taken together, these results demonstrate that chemorepulsion by 8-CPT-cAMP requires the presence of an attractant.

### Guidance uses receptor activity gradients, not attractant gradients

Our findings show that interacting ligands guide cells in unexpected ways. The system’s complexity makes it hard to understand the underlying mechanisms intuitively. We therefore adapted our simulations of chemotaxis (Tweedy et al., 2016, Tweedy et al., 2020) to include steady state competition for receptor sites between ligands. We started without dynamic changes caused by ligand degradation, meaning cells are unable to alter the ligands that have been applied to them, and we can be sure they are responding to the gradients we impose. Simulations of a gradient of competitive antagonist with a background of non-degradable attractant (Fig. 2A) predicted effective negative chemotaxis away from the antagonist. Interestingly, this effect is most pronounced at the bottom of the gradient, and diminishes as the concentration of the repellent rises (Fig. 2B, C). No appreciable repulsion is seen at the upper half of antagonist concentrations.

**Fig. 2:**
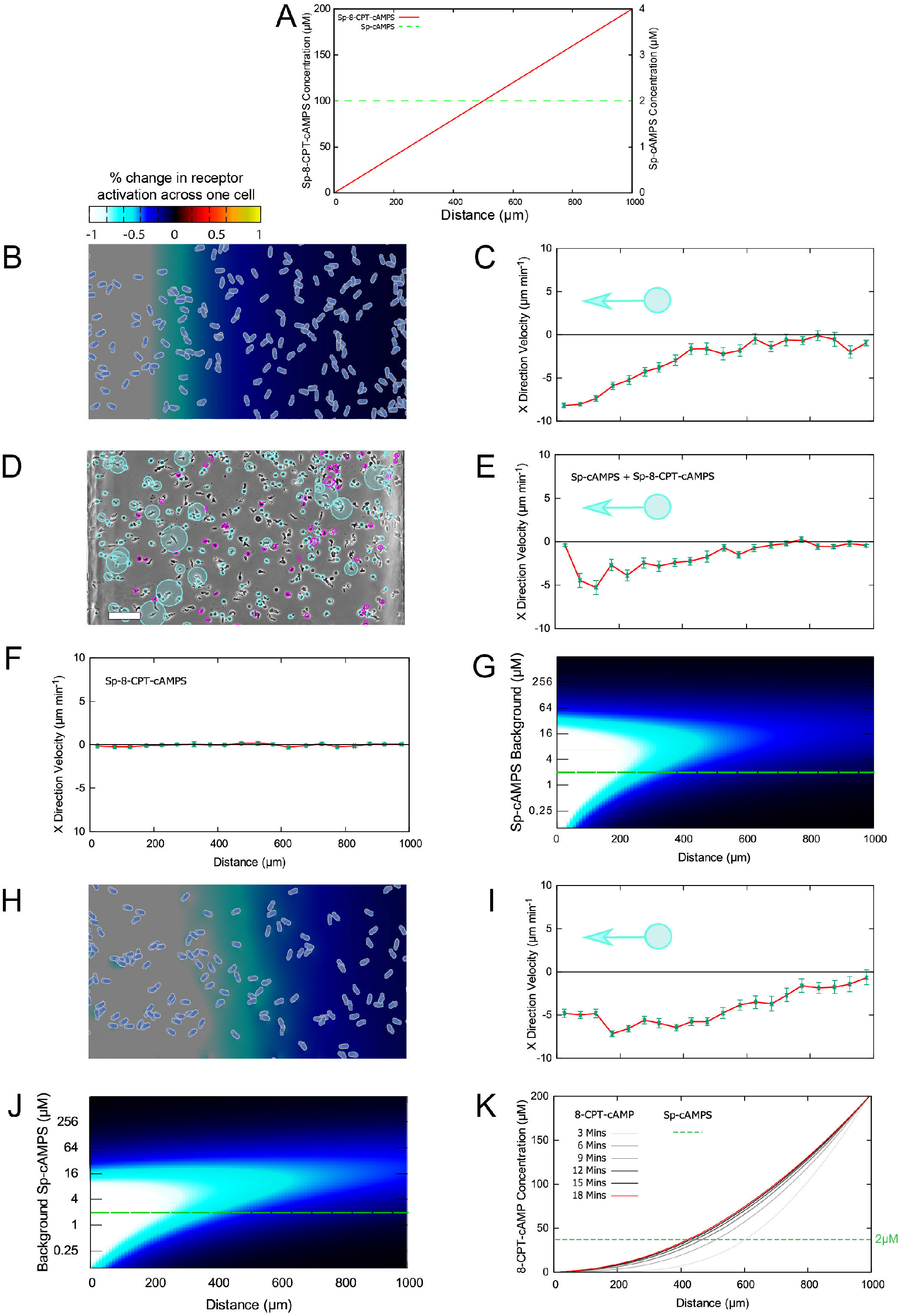
Competitive chemorepulsion using degradable and non-degradable antagonists. **(A)** Concentrations used in this figure. The nondegradable antagonist forms a linear steady state between the leftmost concentration of 0 and the rightmost concentration of 200μM. A background agonist is included. Concentrations are optimised for the antagonist Sp-8-CPT-cAMPS and the agonist Sp-cAMPS **(B**,**C)** Modelled chemotaxis under conditions described in (A). Colours show the local difference in receptor activity, with blue through white showing increasing leftward (i.e. repulsive) cues, and black to red to yellow showing stronger rightward (i.e. attractive) cues. (C) shows quantitation, predicting clear repulsion at the left-hand side of the chamber. **(D, E)** Experimental verification of modelled data. Cell tracker overlay shows magenta circles for NC4 cells moving rightwards and cyan circles for cells moving leftwards. The repellent effect is again strongest at low antagonist concentrations. Scale bar 100μm. **(F)** Velocity of real cells stimulated with Sp-8-CPT-cAMPS gradient alone. Cells barely move in this condition. **(G)** Heat map showing differences in receptor activity, by position in the gradient (x-axis) and background Sp-cAMPS concentration (y-axis), assuming a linear 0-200μM antagonist gradient. Higher concentrations of Sp-cAMPS improve chemorepulsion up to a point, beyond which Sp-cAMPS saturation inhibits guidance as most receptors are activated. The green dotted line indicates the concentration value used in the experiments. **(H, I)** Same as (B, C), but using an antagonist that the cells can degrade locally. This leads to stronger, wider-ranging repulsion. **(J)** As for (G), but for a degradable antagonist. **(K)** Time course showing simulation of the evolution of a degradable antagonist gradient and formation of a near-exponential gradient. Green points in panels C, E, F, I show mean ± SE of technical replicates (n ≥ 53, 23, 49, 40 respectively) for cells at the indicated positions in a representative experiment.

To reproduce this model in real cells, we required a ligand that combined being a full antagonist (such as 8-CPT-cAMP) with resistance to phosphodiesterase (such as Sp-cAMPS). Fortunately these modifications are in different parts of the molecule, meaning that Sp-8-CPT-cAMPS is both an antagonist and nondegradable. We found that experiments using a combination of Sp-8-CPT-cAMPS and Sp-cAMPS reproduce the behaviour of the models almost perfectly (Fig. 2D, E). A gradient of Sp-8-CPT-cAMPS alone gives no appreciable directionality across the whole bridge (Fig 2F), confirming that this molecule is a simple antagonist and an agonist must be present for repulsion to be effective.

For a quantitative understanding of antagonistic repulsion, we built an analytical model of receptor occupancy across the whole of the chemotaxis chamber (See SI for details). We used this to generate guidance heat-maps (Fig. 2G) showing the effect of different background attractant concentrations. This predicts an optimal background concentration of Sp-cAMPS of between 4 and 16μM, about 4-16 times its K_*d*_ for the receptor, cAR1 (Fig. 2G). Below this, responses decline as the background agonist concentration drops, and cell behaviour therefore approaches the unsteered state. Predictions for different concentrations allowed us to refine our conceptual understanding. The agonist and antagonists actively compete for sites. Therefore high agonist concentrations saturate receptor sites, diminishing the capacity of the antagonist to cause spatial bias. Low agonist concentrations are less effective because there is less active signal to outcompete.

The non-degradable inhibitor gave a much less robust response, across a smaller proportion of the bridge, than an inhibitor that can be broken down (see Fig. 1D, G). This was surprising, as we had expected degradation to mute the repellent’s effect. We tested this using simulations, which replicated both the enhanced strength and the range of a degradable chemorepellent (Fig. 2H-K). The simulation revealed that the interaction of diffusion and degradation creates an approximately exponential gradient across the chamber (Fig. 2L), exposing all cells to a steeper gradient of receptor inhibition. Thus cells can obtain more information from chemorepellents by degrading them, just as they can with chemoattractant (Tweedy and Insall, 2020).

### Antagonist gradients can reverse positive cues

We tested whether two gradients could interact together to give repulsion. Simulation of a combination of agonist and antagonist gradients (Fig. 3A-B) predicted a negative response (Fig. 3C-F), as long as the agonist started at a significant level (Fig. 3B). For both attractant alone and the combination, chemotaxis was clearest at the lower end of the gradient (Fig. 3E, F), becoming less effective as the concentrations got higher and the receptors saturated. Real cells behaved exactly as the model predicted (Fig. 3G-J). The dose response of this repulsion is complex - agonist and antagonist gradients in the same direction can be repulsive, but only if the growth in antagonist is enough to overcome the attractive signal (and reverse the gradient of activated cAR1 receptors).

**Fig. 3:**
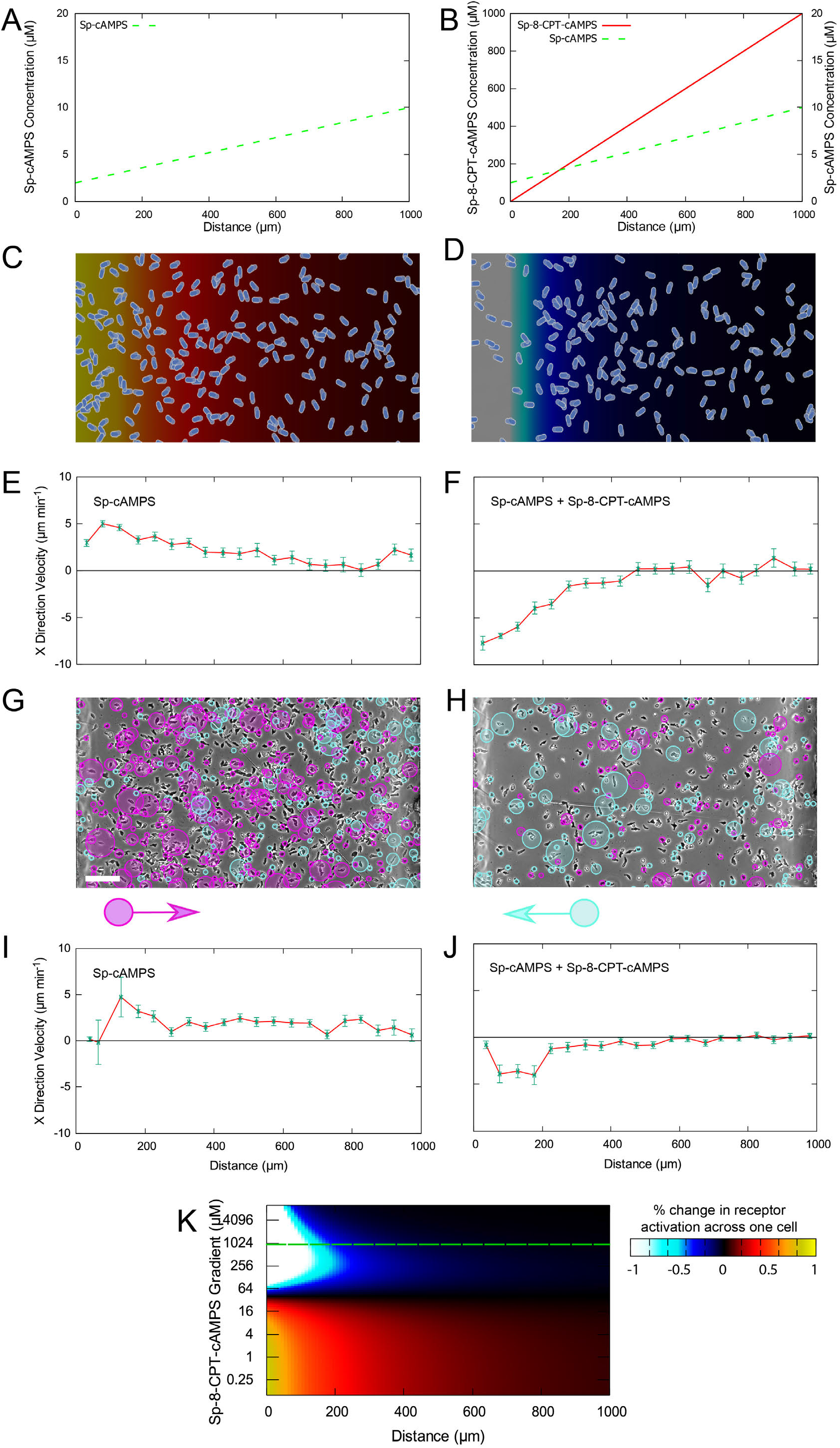
Interference between colinear agonist and antagonist gradients. **(A, B)** Diagrams of the two conditions in the figure, a 2-10μM agonist (Sp-cAMPS) gradient alone (A), and alongside a 0-1mM antagonist (Sp-8-CPT-cAMPS) gradient (B). **(C-F)** Modelled chemotaxis under conditions shown in (A,B). The agonist alone drives positive chemotaxis (C), but overall guidance is repellent when the antagonist is introduced (D). (E, F) show quantitative analysis of rightward velocity. In both cases, guidance is strongest at the bottom end of the gradient toward the left. **(G-J)** Experimental validation of (C-F). Tracking of NC4 cells shows clearly the attractive (G) and repellent (H) regimes, with the rightward velocity component quantified in a representative experiment (I,J). Scale bar 100μm. **(K)** Heat map showing the difference in the proportion of active receptors between the front and the back of each cell, by position in the gradient (x-axis) and steepness of the non-degradable antagonist gradient (y-axis), assuming the presence of the linear agonist gradient shown in (A, B). Shallow antagonist gradients leave the agonist able to guide cells toward the right, but steeper gradients change the direction of guidance. The green dotted line shows the conditions used in the rest of the figure. Green points in panels E, F, I, J show mean ± SE of technical replicates (n ≥ 53, 33, 36, 46) for cells at the indicated positions in a representative experiment.

In conclusion, shallow agonist gradients are able to combine with steeper antagonist gradients to steer cells negatively. Above a critical steepness, at which the two balance out, the guidance direction reverses (Fig. 3K).

### Interactions between degradable and non-degradable attractants and repellents

Since cells can increase the strength and range of negative chemotaxis by locally degrading the antagonist (Fig. 2G, J), we examined the interactions between degradable attractants and repellents in detail. Cells completely degrade low concentrations of cAMP signal, which means they cannot perceive the interfering gradient. However, by adding a small amount of nondegradable Sp-cAMPS, we created an agonist gradient that always had a detectable signal (Fig. 4A). This allowed experiments with a near-exponential gradient across most of the chamber, while still maintaining competition between antagonist and agonist.

**Fig. 4:**
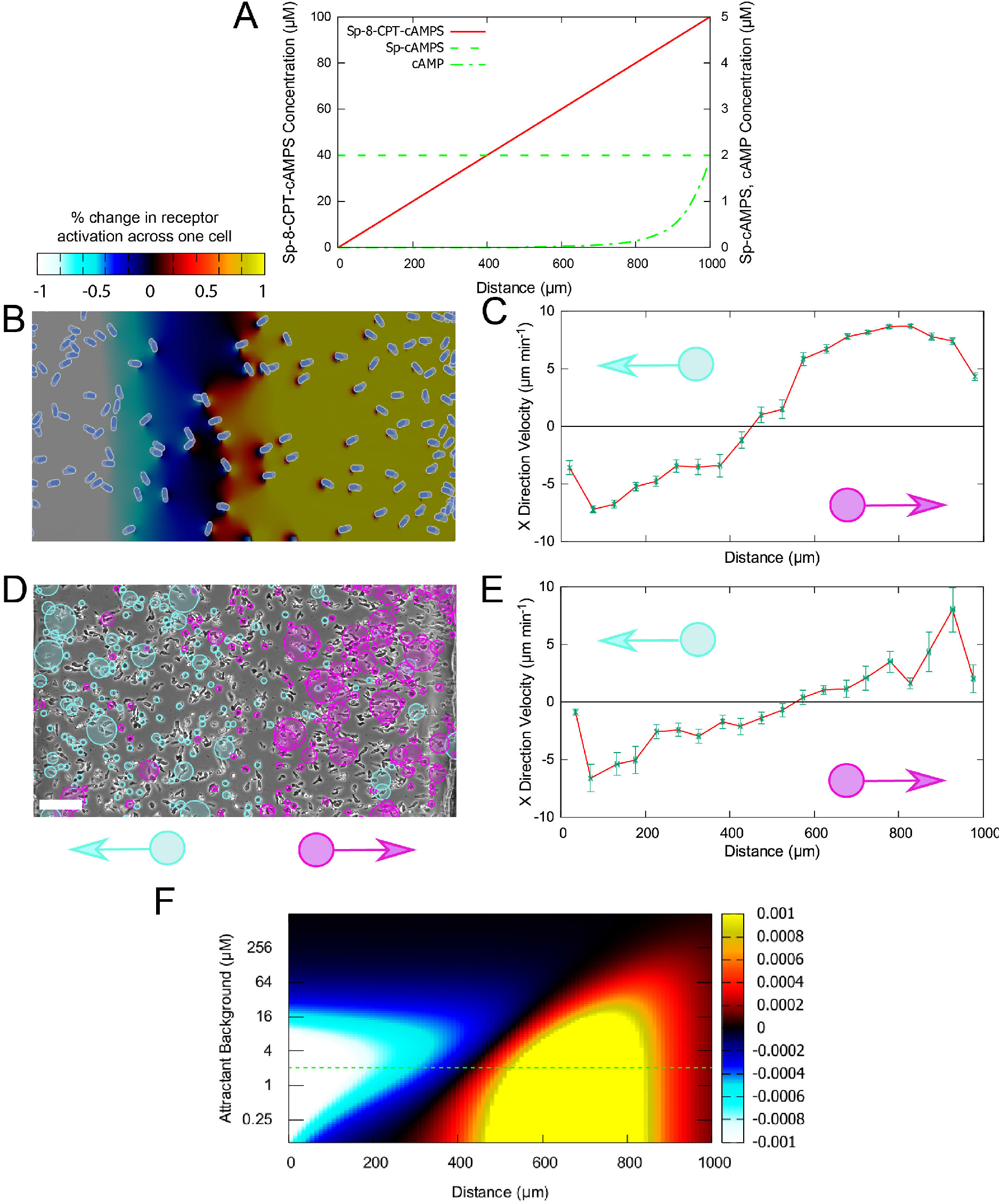
Divergent guidance from 3-ligand interaction with a degradable agonist. **(A)** Diagram of the environmental conditions used. A linear gradient of 0-100μM Sp-8-CPT-cAMPS antagonises a 0-5μM (initially, before degradation) cAMP gradient, which becomes exponential due to attractant degradation. A 2μM background of Sp-cAMPS maintains some receptor activation. **(B)** Modelled chemotaxis, with background colour showing receptor activity difference across the cell. Expected receptor activity has a minimum in the middle of the chamber, leading cells to either side. **(C)** Quantification of the rightward velocity in (B). **(D, E)** Experimental verification of the predictions in (B,C), with NC4 cells moving left highlighted in cyan and cells moving right highlighted in purple. Scale 100μm. **(F)** Heat map of the calculated difference in proportion of receptor activity across cells for the conditions described in (A), with antagonist background concentration varying on the y axis. The black zone in the middle of the map shows the midpoint where cell behaviour diverges. Green points in panels C & E show mean ± SE of technical repeats (n ≥ 23, 32) for cells at the indicated positions of a representative experiment (E) or 3 repeated simulations (C).

Simulations of this setup made a striking prediction – the cells diverged at the midpoint of the chamber, with those in the left half moving left, those on the right half moving right (Fig. 4B, C), opening up a gap in the centre. Guidance cues were very strong at both ends (Fig. 4C). Real cells exactly replicated this behaviour (Fig. 4D, E; Movie 3), again showing the accuracy of the simulations. Plotting simulated receptor occupancy shows that this combination allows an exponential gradient of cAMP to dominate the right half of the chamber (Fig. 4F), climbing far faster than the antagonist gradient. In contrast, the cAMP was at near zero levels on the left-hand side of the chamber, so the gradient was too shallow to affect the repellent effect of the other ligands.

This experimental condition - in which the cells change direction depending on their position in the gradient – is very unusual. Cells chemotax differently depending on their position (Fig. 4F). In a simple attractant gradient, the guidance cues are stronger at the low end, but at higher concentrations the difference between front and rear becomes less and they lose their sense of direction (for example Fig. 3G, I). However, to our knowledge it has not been seen before. The behavioural turning point is neither a source of attractant nor of antagonist, but the result of a complex competition between different ligands.

We also tested whether the opposite was true, and degradable antagonists could lead to convergence in the middle of the gradient. Simulations predicted that degradable antagonist gradients could be shallow enough at their lower ends to be outcompeted by a moderate, linear, agonist gradient. At the same time, higher concentrations and the exponential nature of a degradable antagonist at the high end can overwhelm the linear agonist gradient (Fig. 5A). This leads to a striking maximum of receptor activity in the middle of the chamber, which causes cells from everywhere in the bridge to converge (Fig. 5B, C; movie 4). Experiments with real cells again followed the simulations remarkably accurately, verifying that the model’s prediction was correct (Fig. 5D, E). One difference was extremely interesting and informative - real cells in antagonist ceased making protrusions and stopped moving, rather than continuing with the random movement as we see in unstimulated cells (see movie 4). The principal result is that competition between ligands can direct cells to positions that are different from the source of either ligand, which depends on the K_*d*_ of the antagonist relative to the agonist (Fig. 5F).

**Fig. 5:**
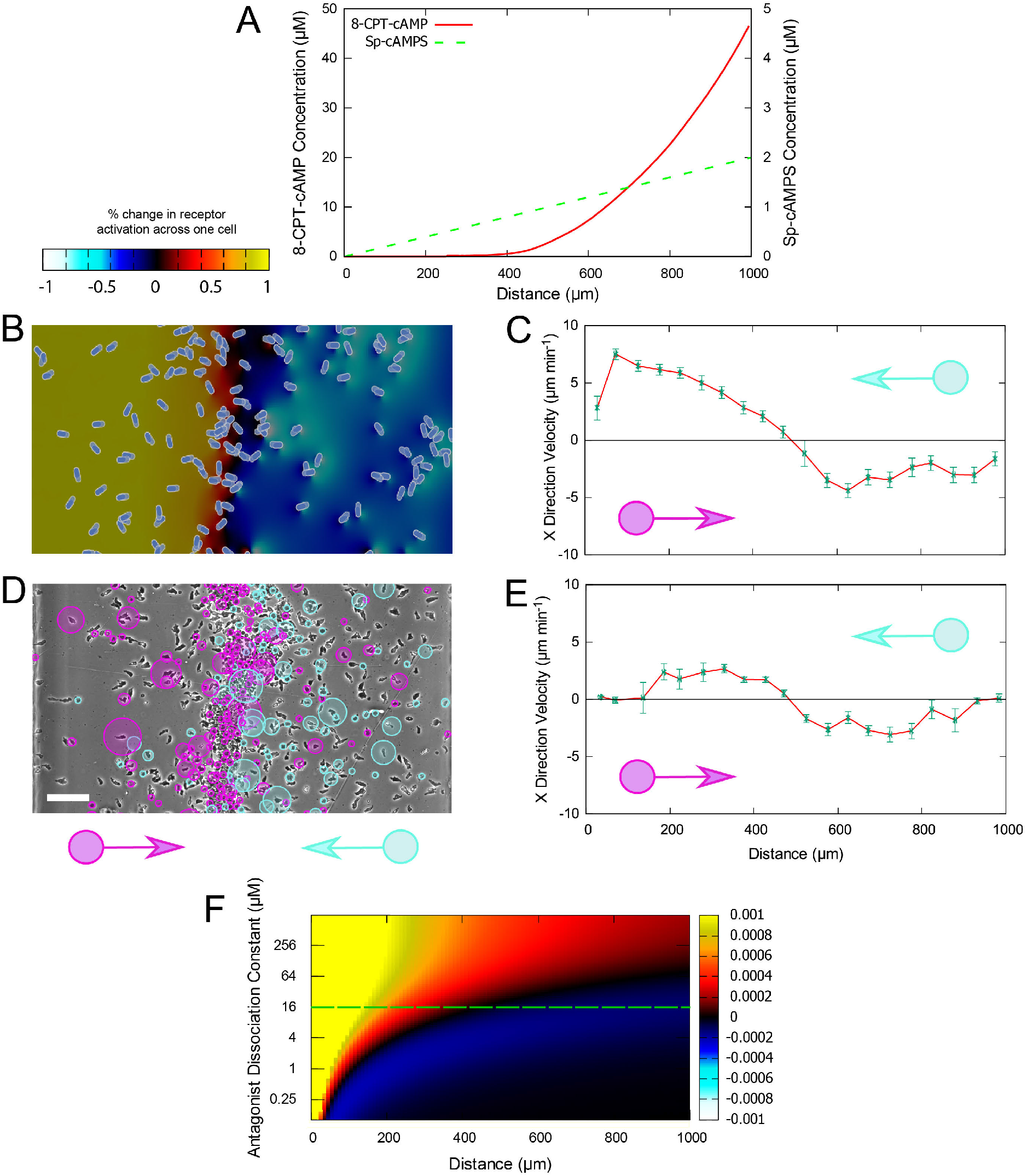
Convergent guidance from stable agonist and degradable antagonist interference. **(A)** Diagram of the environmental conditions used. A linear gradient of 0-2μM Sp-cAMPS provides an agonist signal against which a 0-80μM 8-CPT-cAMP gradient interferes. Cell degradation of the latter makes it reach an exponential shape at equilibrium. **(B)** Modelled chemotaxis, with background colour showing receptor activity difference across the cell. Cells converge on a central point, rather than being drawn to either well. **(C)** Quantification of the rightward velocity in (B). **(D, E)** Experimental verification of the predictions in (B,C), with NC4 cells moving left highlighted in cyan and cells moving right highlighted in purple. Scale bar = 100μm. **(F)** Heat map of the calculated receptor activity gradient for the conditions described in (A), with antagonist Kd varying on the y axis. The black zone in the middle of the map shows the midpoint where cell behaviour converges. Green points in panels C & E show mean ± SE (n ≥ 23, 13) for cells at the indicated positions, in 3 experiments (C) or representative experiment (E).

### Complex outcomes from mixed signals: Chemorepulsion by competing gradients

Many physiologically important examples of chemotaxis are remarkably complex. For example, there are more than 40 chemokines, which are transduced by around 20 receptors; each receptor may detect many chemokines, and some chemokines activate different receptors to different degrees. To explore how this complexity affects chemotaxis, we tested the effect on chemotaxis of mixing different ligands to the same receptor.

One feature of receptor behaviour is particularly important. Receptors are normally in an ‘inactive’ state. When they are in an ‘active’ state, they bind to G-proteins and cause effects inside the cell. When a perfect receptor binds to its ligand, it always changes from inactive to active. However, real-life receptors are not perfect; ligand binding only activates a fraction. This is termed the efficacy, and it differentiates an agonist and an antagonist. cAMP, which is a nearly perfect agonist, has an efficacy close to 1, whereas the antagonist 8-CPT-cAMP has an efficacy of effectively zero. The nondegradable analogue Sp-cAMPS is also very efficient. Ligands with intermediate efficacies are called partial agonists. They appear to be weaker attractants, but because they still activate some receptors they can still generate positive guidance cues. This is true of the cAMP analogue Rp-cAMPS.

We simulated three conditions: a gradient of agonist, a gradient of partial agonist, and both gradients expressed together (Fig 6A-I). Cells chemotax up simple gradients of either the partial agonist Rp-cAMPS (Fig. 6D,G) or the efficient agonist Sp-cAMPS (Fig. 6E,H). When these gradients are superimposed, the simulations make an extraordinary prediction. Not only do the two gradients not reinforce one another, but together they will actively repel cells (Fig. 6F, I, J).

**Fig. 6:**
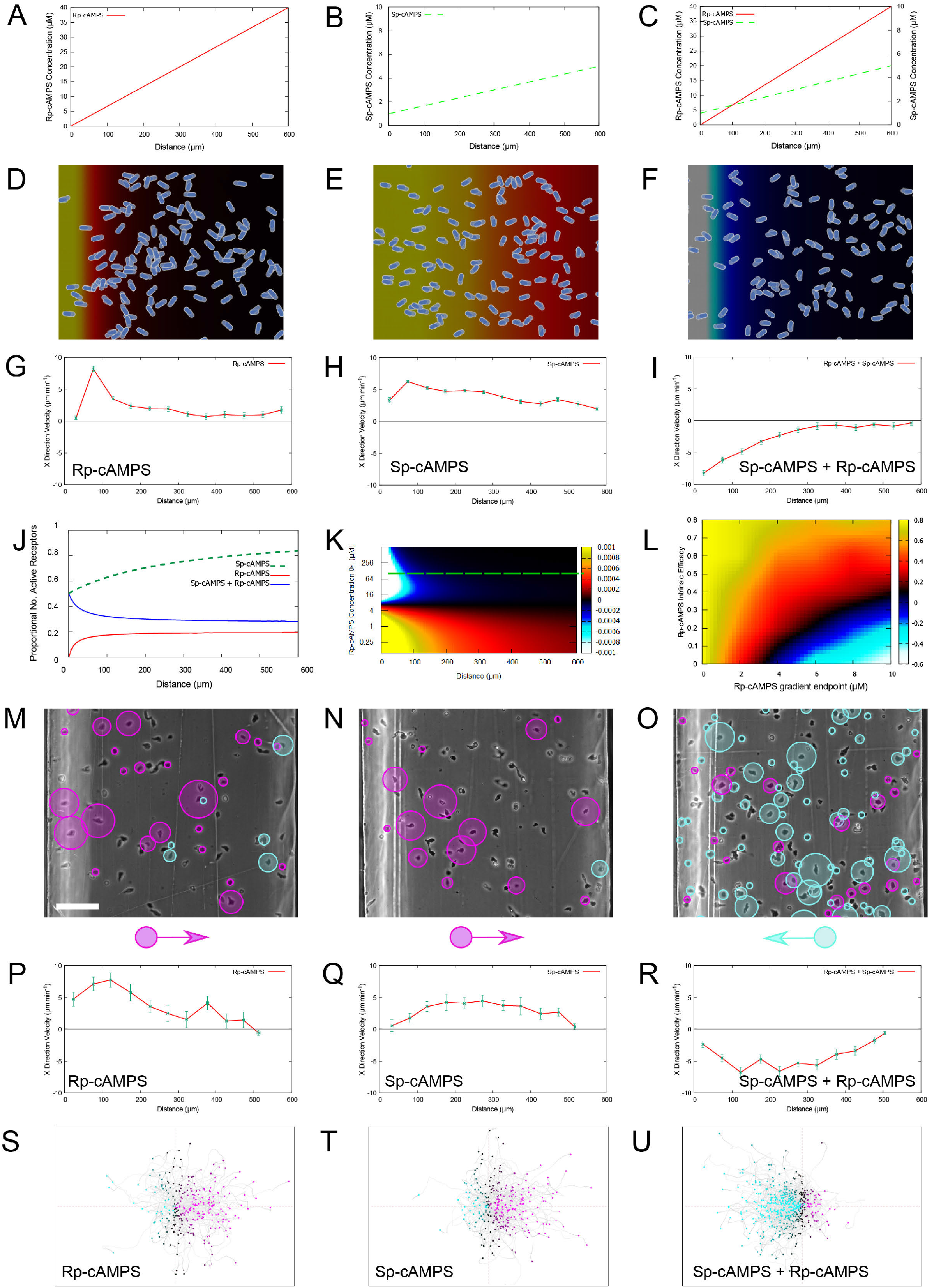
Two aligned attractant gradients may cause repulsion. **(A-C)** Diagrams of the conditions in the figure, a 0-40μM gradient of the partial agonist Rp-cAMPS (A), a 1-5μM gradient of the agonist (Sp-cAMPS) (B), and both together (C). **(D-F)** Modelled chemotaxis in conditions (A-C). Both the partial agonist (D) and the agonist (E) drive positive chemotaxis, but cells are repelled when both gradients are present (F). **(G-I)** Rightward velocity quantified for (D-F). In all three cases, guidance is strongest at the bottom end of the gradient(s) toward the left. **(J)** Computed fraction of receptors that are activated across the bridge of the chamber, under each of the conditions **(K)** Heat map of the receptor activity differences across the chamber when varying the steepness of the partial agonist gradient, assuming its efficacy is 0.3 and presence of the linear agonist gradient shown in (B, C). Shallow partial agonist gradients allow the full agonist to drive positive chemotaxis, but steeper gradients switch the cues to repellent ones. The green dotted line shows the predicted behaviour for the gradient used in the rest of the figure. **(L)** Heat map showing the proportion of simulated cells that show positive (warm colours) or negative (cold colours) chemotaxis, as the intrinsic efficacy and final concentration of the imperfect agonist are varied. The linear attractant gradient illustrated in panel B competes against the linear imperfect inhibitor. Measurements were made over 40min of simulated chemotaxis. **(M-U)** Experimental validation of (D-I). Tracked *aca*A^-^ cells show moderate attraction for both Rp-cAMPS and Sp-cAMPS individually (M, N), and repulsion when both are present (O). Scale bar = 100μm. Rightward velocity components for (P-R) and spider plots showing the paths of tracked cells (S-U, upgradient paths in magena, down-gradient in cyan) are shown below them. Green points in panels G, H, I, M, N, O show mean ± SE (n ≥ 68, 80, 48, 8, 3, 19) for cells at the indicated positions, in 3 simulations (G-I) or a representative experiment (M-O).

We wanted to know if it could be possible for a more efficient agonist to reinforce a chemoattractant signal. Mathematical analysis indicates that a second competing agonist cannot reinforce signals if the first agonist gradient is steep. The switch to repulsive behaviour we witnessed requires two things - a background level of the first attractant throughout the gradients, and lower efficacy for the second attractant (Fig. 6L shows the effects of different efficacies - we estimate the efficacy of Rp-cAMPS to be around 0.3, meaning that of 10 receptors bound to Rp-cAMPS, 3 will become activated). We examined the results we would expect from different gradient steepnesses of a partial agonist with this efficacy (Fig. 6K). Shallower gradients leave the full agonist able to dominate. Above a critical steepness, at which the two signals cancel one another out (around 7μM, in this case) guidance cues become repellent.

We tested our predictions with real cells. Experiments accurately replicated the remarkable result (Fig. 6M-R; Movie 5) - two gradients, each attractive on their own, become repulsive when combined. Thus mathematical modelling is accurately able to predict how competition of ligands for the same receptors can give negative chemotaxis.

## Discussion

Chemotaxis is fundamentally important to normal eukaryotic biology, and thus widely studied. Many important cases - for example, neutrophils responding to infections by chemotaxing towards chemokines - involve many different ligands competing for a smaller pool of receptors. For example, the CXCR2 receptor binds to CXCLs 1, 2, 3, 5 and 8; all are physiologically important(Tharp et al., 2006). These ligands’ chemotactic effects are generally studied individually; we do not know of experiments studying chemotaxis combinations of different CXCLs. This, combined with the need for mathematical modelling to dissect the interactions, means that the role of competition in negative chemotaxis has not been seen previously. The data we have described show that this competition is fundamentally important to chemotaxis in vivo, and understanding it will yield important - if counterintuitive - results.

We have shown how GPCRs can mediate negative chemotaxis without invoking any currently unknown molecular mechanisms. As this relies on competition between ligands, further study into systems with apparent GPCR-mediated chemorepulsion will need to account for competing signals that communicate through the same receptor. Crucially, the findings are not limited to GPCRs: the same mechanisms will work for any receptor type. For example, Robo-mediated guidance is known to involve both full length Slit and Slit-N(Bhat, 2017), suggesting competitive rather than direct repulsion, and the same is true for the pairing of semaphorins and Slit-C with Plexin-1A(Delloye-Bourgeois et al., 2015).

Whilst systems with only one or two known signals might become straightforward when viewed in the context of this work, others are more complex. The chemokine system, for example, involves complex, promiscuous interactions between receptors and ligands, and an array of ligand modifications that change the K_*d*_ and efficacy for different receptor-ligand interactions. This plethora of parts has been described as ‘redundancy’(Wolf et al., 2008), but our finding that competition between ligands gives complex effects (Fig 6) disagrees. It is more likely that a healthy immune system places chemokines at a series of tipping points, ready to mediate attraction or repulsion in specific cell types at different times and places. This would certainly explain the high failure rate of chemokine receptor targeting drugs; understanding it would improve development of new ones.

This also emphasises the importance of studying simplified systems. Our studies succeeded because we could eliminate breakdown of attractants and repellents by cells (using Sp-modifications), and prevent them from secreting their own attractants (using caffeine and by knocking out adenylyl cyclase). Without this, even Dictyostelium is too complex to understand quantitatively, even though it essentially responds to only one chemoattractant. Immune cells chemotax to multiple signals, using many receptors, many of which (for example LTB4(Lämmermann et al., 2013)) are autocrine.

More generally, seeing chemotaxis from the cell’s view (as a simple spatial bias in receptor activation), rather than the experimenter’s (multiple, separate attractant gradients), introduces mechanisms of control that apply in any context. Our investigation into 8-CPT-cAMP shows that the strength of an effective repellent is tuned by changes in the concentration of a ubiquitous agonist, and turned off entirely if the agonist is not present. We show that the inclusion of a competitive antagonist can turn an attractive guidance cue into a repellent one (Fig 3). This would be an excellent mechanism for resolving inflammation. After some threshold is passed, an attractive signal could be competed out by a different agonist, reversing the directional instruction.

Finally, we repeatedly see a motif of strong guidance at the low end of a gradient, whether for positive or negative chemotaxis. This is because the model assumes that current receptor activity is the sole source of information for the cell. However this information is processed, it is important to remember that cells go where receptor and G-protein activation tell them to.

We have shown many ways in which chemotaxis is complex and counterintuitive. Attractant degradation (Tweedy et al., 2016) can be vital to long range guidance, and that local topology can interact with guidance cues, even as far as to lead cells away from major attractant sources (Tweedy et al., 2020). We now add ligand competition to the complex mechanisms that guide cells. Our examples of ligand competition use effectively one-dimensional environments - the cell’s chemotactic stimuli are defined simply by how far across the chamber they are. If we extend this into 2 and 3 dimensions, competition will be even more dynamic and unexpected. Understanding this will require extensive mathematical and computational modelling, combined iteratively with experimental measurements. The counterintuitive nature of these questions means that we could not design informative experiments without computational simulations to test the implications of our current understanding. In any case, it is increasingly clear that real guidance cues emerge from the interactions of several signals, and they must be considered together to be understood.

## Methods

### Cell Culture

Two *Dictyostelium* strains were utilised in this paper, wild type NC4 (WT) and NC4 adenylyl cyclase knockouts (*aca*A^-^). Both were grown on *Klebsiella aerogenes*, passaged once weekly. The *K. aerogenes* was also passaged once weekly. Fresh stocks of bacteria and both NC4 strains were retrieved from −80°C once per month.

### Assay Preparation

For both WT and *aca*A^-^strains, experiments were set up two days in advance by growing cells on four different agar plates - with *K. aerogenes* - at varying cell densities. On the day of the experiment the plate with the most consumed bacteria and least sign of further starvation was harvested, and the cells washed in KK2 buffer (K phosphate pH6.2), with added 2mM MgCl_2_ and 0.1mM CaCl_2_ (to give KK2MC) three times at 1300 rpm for 3 minutes. For WT cells, the cell suspension was adjusted to a cell density of ∼3×10^6^ cells/ml and 4ml was then placed on a series of 60mm plates with 1% agar. After a couple of minutes any excess liquid was drained and the cells were then left to starve for approximately 5 hours.

Agar plates were left in the refrigerator if a longer development time – perhaps for testing subsequent conditions – was required. At the onset of streaming plates were harvested, the cells washed and resuspended with 2mM caffeine - to supress further cAMP secretion – at ∼6×10^5^ cells/ml. Cells were then left to settle on 22mm^2^ slides, using 200μl suspension, for 20 minutes before being used in an Insall chamber assay.

For *aca*A^-^ cells, the cell suspension was adjusted to a cell density of ∼1×10^7^ cells/ml and 10ml was then placed into a small flask. As these cells do not secrete cAMP they cannot develop via the usual method of starvation induced cAMP relays; instead they must instead be treated with external pulses of cAMP. The suspension was mixed on a rotating table for 1 hour, so the cells consume any residual bacteria and begin to starve, then treated with 300nM cAMP every 6 minutes for 4 hours, while still being mixed. Cells were then washed and resuspended – without caffeine – at ∼6×10^5^ cells/ml. Cells were then left to settle on slides as with WT above, before use in an Insall chamber.

### Chemotaxis Chamber Assays

Insall chambers & their use are described in ref. (Muinonen-Martin et al., 2010). In all uses, buffer containing the chemical conditions required for the inner well was initially placed everywhere. The coverslip with cells adhered was then placed over the chamber, keeping the tips of the outer well free. The buffer in the outside well was replaced by new buffer containing the chemical conditions required for the outer well. If using WT cells the chemical condition containing buffers also included 2mM caffeine. The chamber was then left to equilibrate for around 15 minutes before filming the cells on the bridge of the chamber using a 10x objective in a Nikon Ti2 timelapse microscope with a Retiga R6 digital camera for around 20 minutes. This was sufficient time to obtain data on the directional bias at different point along the gradient at equilibrium.

A detailed protocol can be found in (Muinonen-Martin et al., 2013).

### Micropipette Assays

For this assay only WT cells were used; they were prepared as for chemotaxis chambers until the developing cells were at the onset of streaming, then harvested, washed and resuspended in 2ml of buffer. Enough suspension to give an appropriately low cell density was placed in a 35mm glass bottom dish containing no caffeine, caffeine or caffeine plus 2μM Sp-cAMPS. The micro-pipette (Eppendorf Nanotip Gold) was loaded with 10mM 8-CPT-cAMP, pressurized and lowered gently into the suspension. Images were recorded on an Olympus microscope at 10x magnification.

### Data Analysis

Data from live cell imaging was analysed using an ImageJ plugin written by L. Tweedy(Tweedy et al., 2016), giving positional data of cells at different time points. This data were converted into an x-direction velocity and average position for each tracked cell, such that an average velocity and error on the velocity could be found for each cell occupying a defined partition of the bridge.

Due to inter-experiment variability, local cell speeds or velocities were plotted as mean ±SE of cells at the indicated location in a single representative experiment; all experiments were repeated at least three times on different days, with consistent results.

### Simulations

Simulations run are based on a Java code written by L. Tweedy(Tweedy et al., 2016), modified according to the needs of the project. Modifications include: inclusion of up to three competing chemicals of varying intrinsic efficacies, a rendering scheme illustrating the rate of change receptor activation with space and the possibility of accounting for chemical degradation in the wells due to cells located there.

### Wild-Type NC4 Bridge Chamber Assays

Cells were starved on agar for approximately 4 hours, examined, and harvested when centres of cell aggregation were visible across the plate, but before cells had started to stream. At this time the cells were harvested from a plate using 1ml KK2, washed into 1ml KK2MC, and diluted 15x into another 1ml KK2MC + 2mM caffeine. After mixing thoroughly once more, 200μl suspension was pipetted onto 22mm^2^ cover slips and allowed to settle for at least 20 minutes before assembling the chamber. This was then left for about 10 minutes for the gradients to equilibrate before filming the 1mm bridge using a 10x objective in a Nikon Ti2 timelapse microscope with a Retiga R6 digital camera for around 20 minutes. This was sufficient time to obtain data on the directional bias at different point along the gradient at equilibrium.

## Acknowledgements

We are grateful to Cancer Research UK (CRUK) for CRUK core funding to the CRUK Beatson Institute (A31287) and to RHI (A19257), and to the Wellcome Trust for grant 221786/Z/20/Z to RHI.

## Author contribution

Conceptualisation: AD, LT & RHI. Experimental design: AD & PT. Experimental investigation: AD & PP. Simulations: AD & LT. Writing: AD, LT & RHI. Supervision: LT, PT & RHI. Funding Acquisition: RHI.

## Movies

Movie 1: Micropipette assay: cells responding to 8-CPT-cAMP, then to 8-CPT-cAMP plus Sp-cAMPS

Movie 2: Chamber assay: cells responding to and 8-CPT-cAMP gradient, homogeneous Sp-cAMPS, then both together

Movie 3: Degradable attractant, nondegradable repellent - cells responding to combined Sp8-CPT-cAMPS and cAMP gradient. A linear gradient of 0-100μM Sp-8-CPT-cAMPS antagonises a 0-2μM cAMP gradient, which becomes exponential due to attractant degradation. A 2μM background of Sp-cAMPS maintains some receptor activation.

Movie 4: Degradable repellent, nondegradable attractant - cells responding to combined 8-CPT-cAMP and Sp-cAMPS gradient. A linear gradient of 0-100μM 8-CPT-cAMP, which becomes exponential due to attractant degradation, antagonises a 0-2μM Sp-cAMPS gradient.

Movie 5: Combining two attractant gradients creates a repulsive gradient. Cells responsing to gradients of (1) Sp-cAMPS, (2) Rp-cAMPS, (3) Sp-cAMPS and Rp-cAMPS combined.

